## Supplementary material for "PoreVision: A Program for Enhancing Efficiency and Accuracy in SEM Pore Analyses": ImageJ Protocol.pdf

### Setup

| | | |
|---|---|---|
| 1. | Open ImageJ | 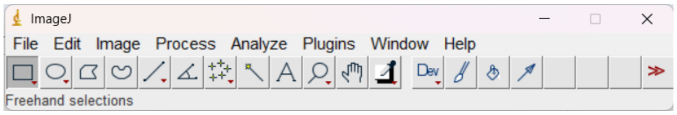 |
| 2. | Click ‘File’ → ‘Open’ | 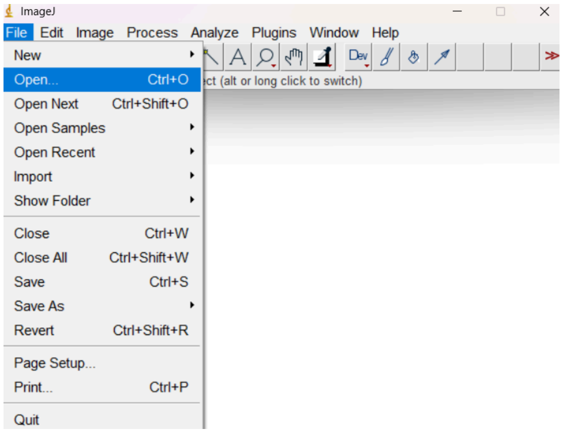 |
| 3. | Select cryogel image | 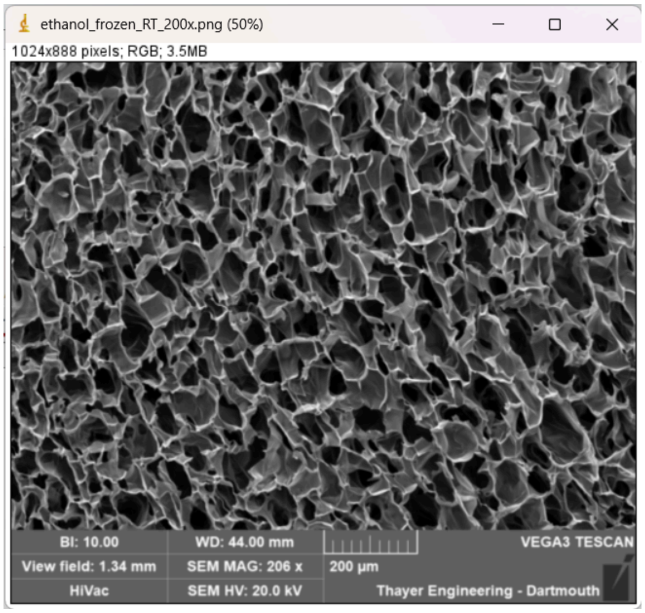 |

| | | |
|---|---|---|
| 4. | Zoom in so that the 200 um scale bar is as large as possible on your screen<br><br>Use the hand tool to move the picture if needed | 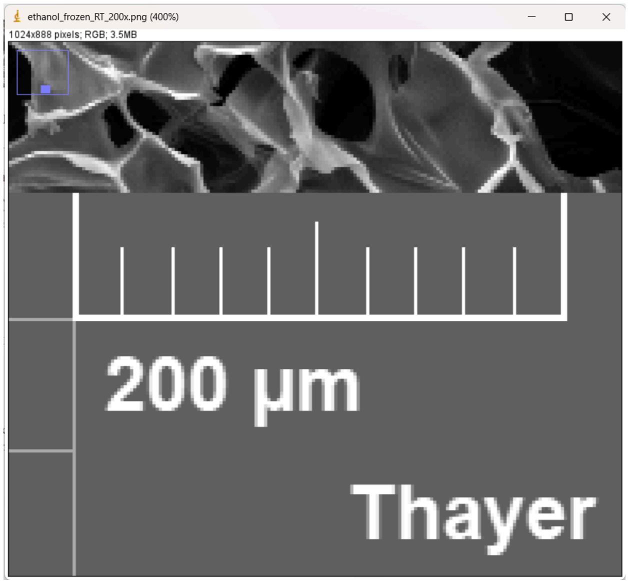 |
| 5. | Click the line tool<br>5th from the left | 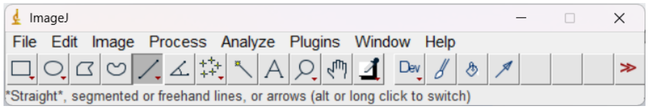 |
| 6. | Click and hold 'shift' to draw a straight line on top of the scale bar | 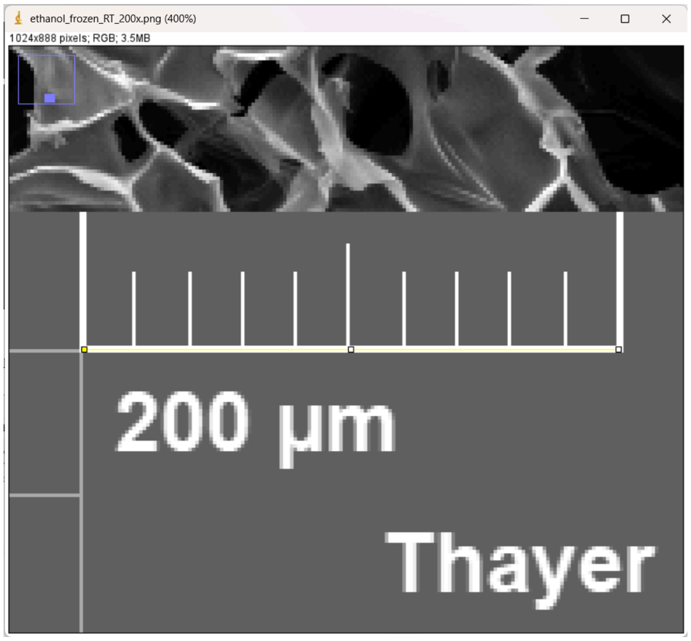 |

<table>
<tr><td>7.</td><td>Click ‘Analyze’ → ‘Set Scale’</td><td>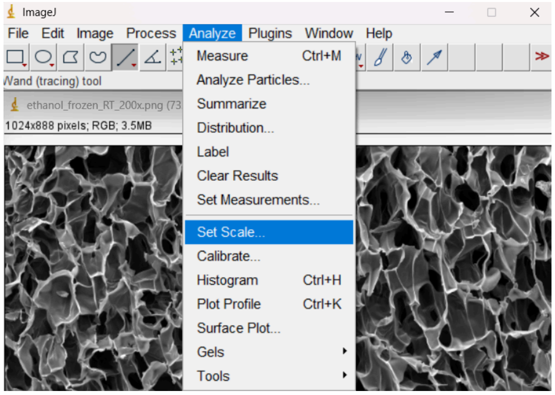
</td></tr>
<tr><td>8.</td><td>In the Set Scale window:<br>

- ‘Distance in pixels’ should automatically populate and be about 145-165 pixels
- Set ‘Known distance’ to 200
- In ‘Unit of length’ type ‘micron’
- ‘Scale:’ should now read about 0.725-0.825 pixels/micron

There might be variation from person to person, that’s okay!

Click ‘OK’</td><td>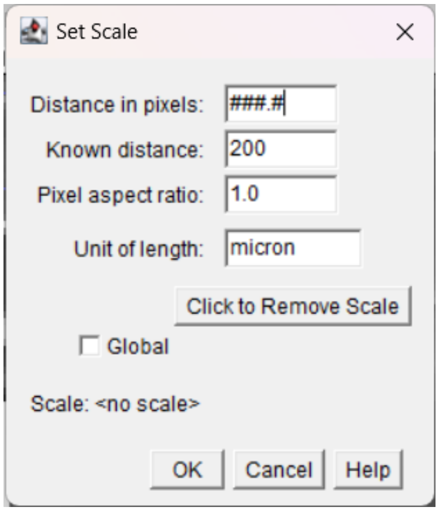
</td></tr>
<tr><td>11.</td><td>Click ‘Analyze’ → ‘Set Measurements…’</td><td>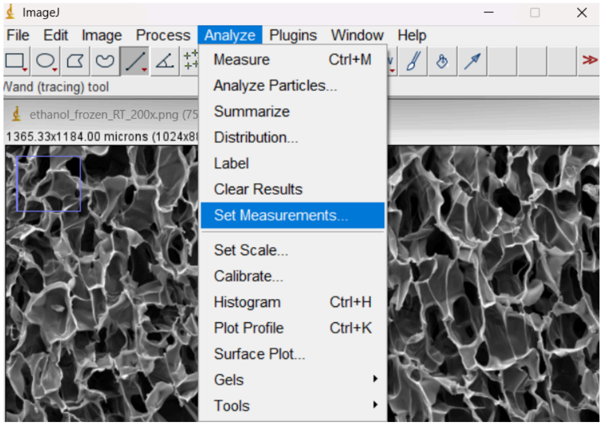
</td></tr>
</table>

| | | |
|---|---|---|
| 12. | Tick the following boxes:<br>● ‘Area’<br>● ‘Skewness’<br>● ‘Perimeter’<br>● ‘Feret’s diameter’<br><br>Click ‘OK’ | 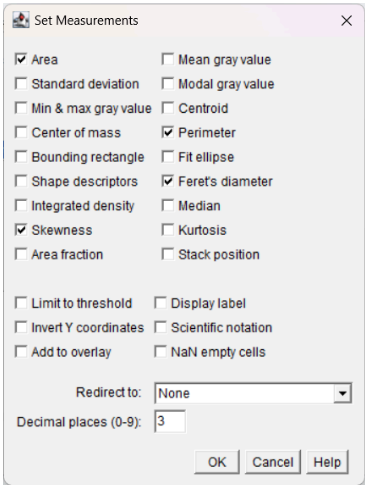 |
| 14. | Click ‘Image’ → ‘Overlay’ → ‘Labels…’ | 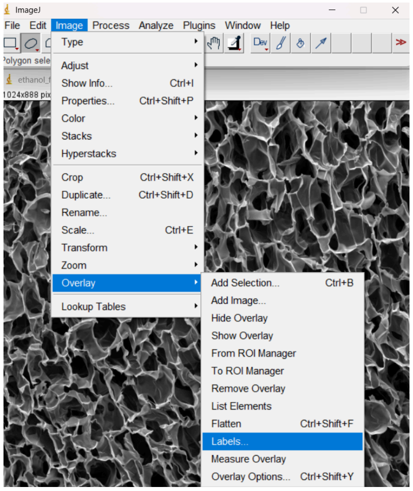 |
| 16. | Change ‘Color’ to ‘yellow’<br><br>Tick the following boxes:<br>● ‘Show labels’<br>● ‘Draw backgrounds’<br>● ‘Bold’<br><br>Click ‘OK’ | 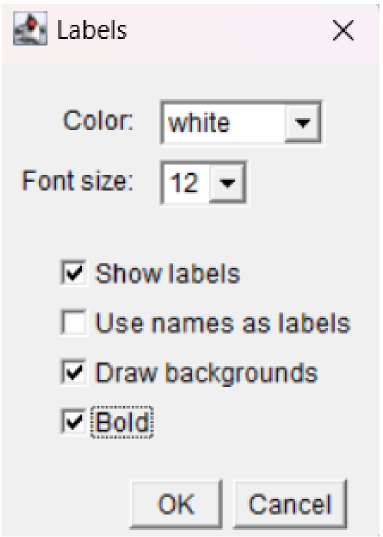 |

### Measurements

| | | |
|---|---|---|
| 1. | Zoom into the top left quadrant<br><br>A blue box in the top left corner should indicate you are zoomed into a quarter of the image | 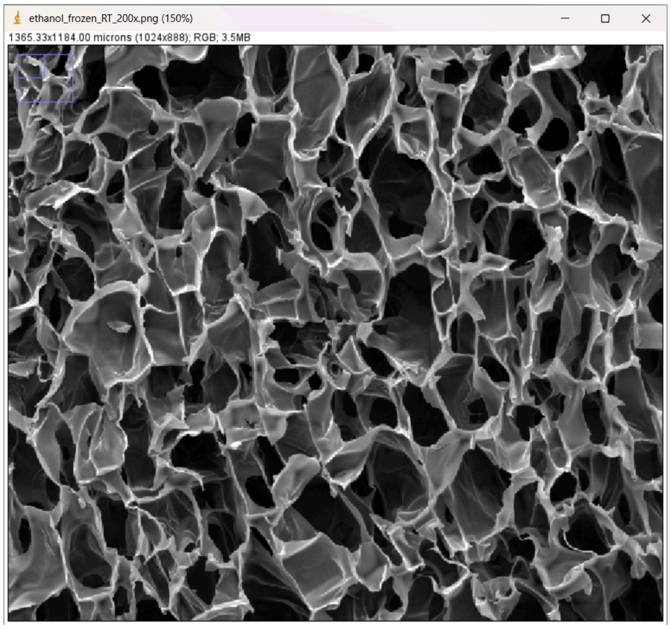<br>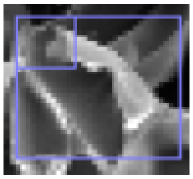 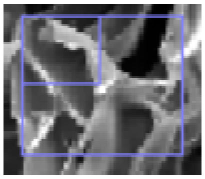 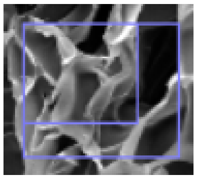<br>Less than 1/4 1/4 More than 1/4 |
| 2. | Long press the ellipse tool (2nd from the left)Click 'Elliptical selections' | 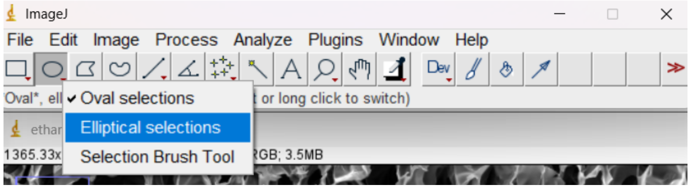 |

<table>
<tr>
<td>3.</td>
<td>Find a pore and create an ellipse around it<br><br>a. Use the nodes to increase or decrease the size and to rotate the ellipse<br><br>b. Try to form the ellipse so it roughly follows the perimeter of the pore<br><br>c. For pores that aren’t easy to fit into an ellipse, use your best judgment<br>(that’s part of the fun!)</td>
<td>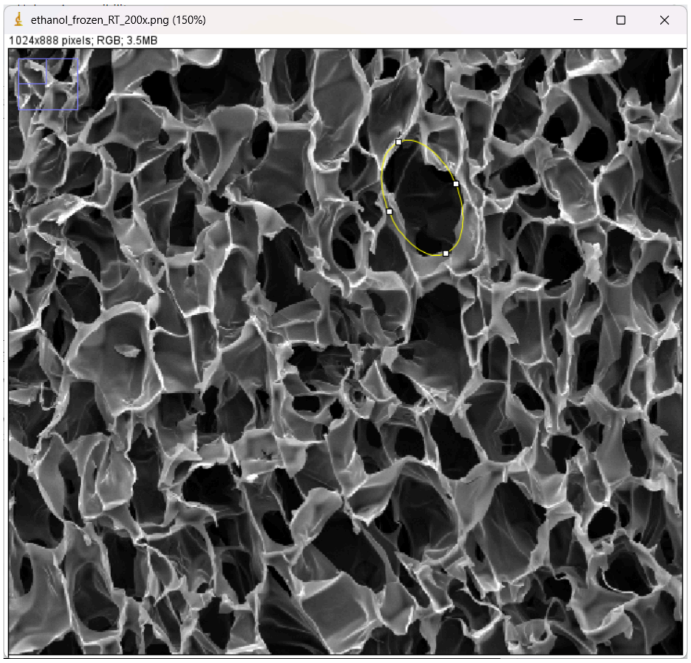
</td>
</tr>
<tr>
<td>4.</td>
<td>After you are satisfied with the ellipse:<br>● On your keyboard press ‘B’ (creates an imprint)<br>● On your keyboard press ‘M’ (adds the measurement to the list)</td>
<td>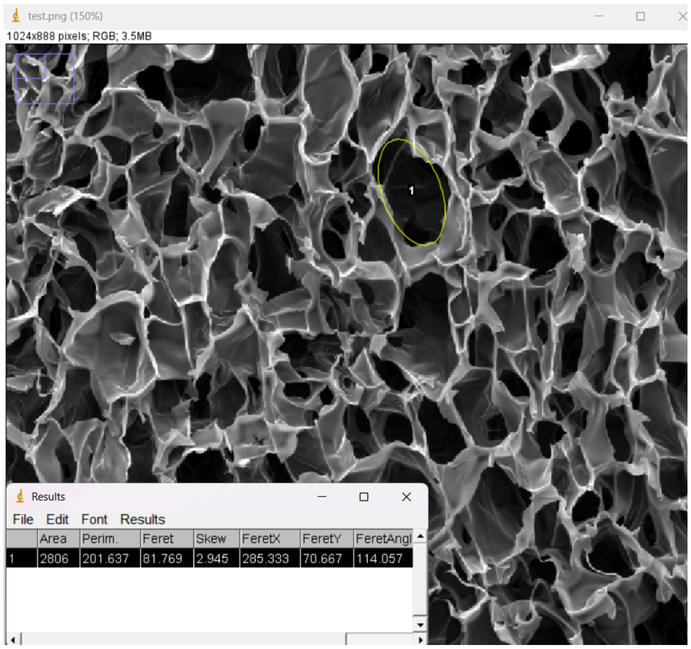
</td>
</tr>
</table>

<table>
<tr>
<td>5.</td>
<td>If you are not satisfied with the ellipse:<br>a. Click the row in ‘Results’ and hit delete to delete the data<br>b. Click the number in the middle of the ellipse to select the ellipse<br>c. Edit!<br>d. Hit ‘B’ on your keyboard<br>e. Hit ‘M’on your keyboard</td>
<td>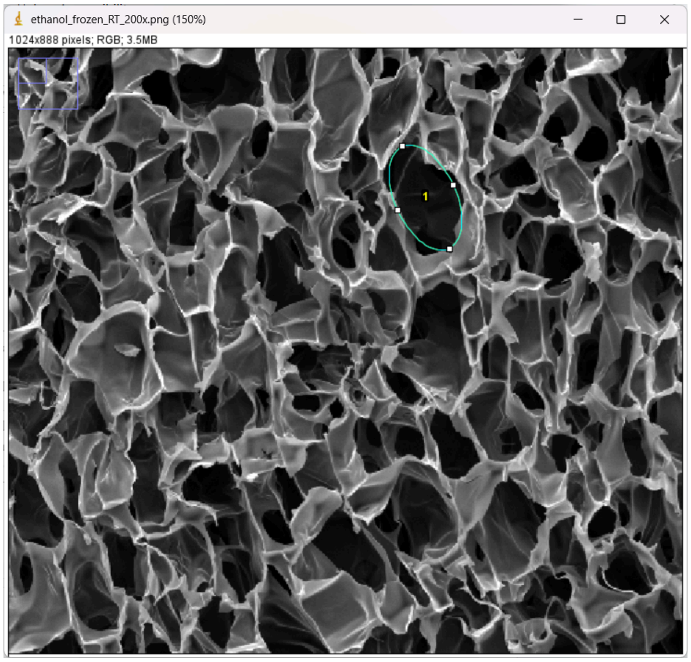
</td>
</tr>
<tr>
<td>6.</td>
<td>Continue this process until there are 15 ellipses in this quadrant<br><br>a. Aim to get a variety of ellipses (large, small, fat, skinny, etc.)<br><br>b. There should also be 15 rows in the ‘Results’ window</td>
<td>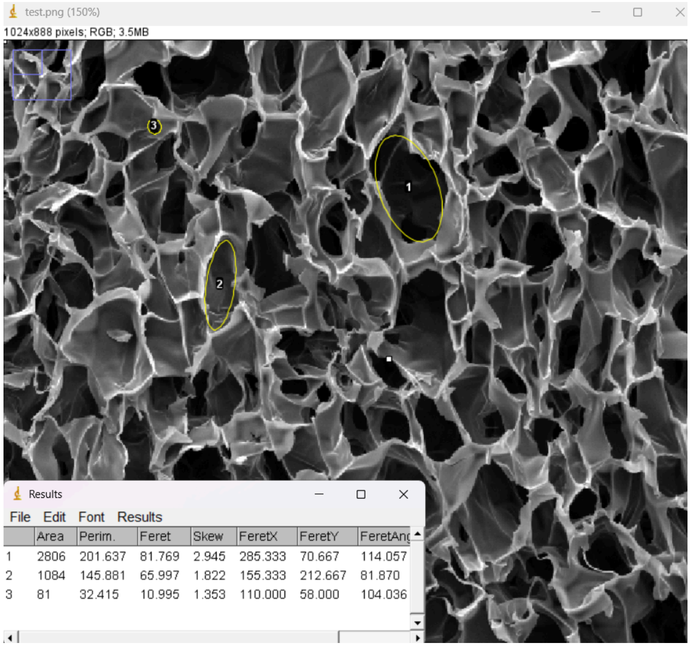
</td>
</tr>
</table>

| | | |
|---|---|---|
| 7. | Repeat steps 4 through 7 for the other three quadrants<br><br>You should have 60 pores outlined and 60 rows of measurements in the ‘Results’ window | 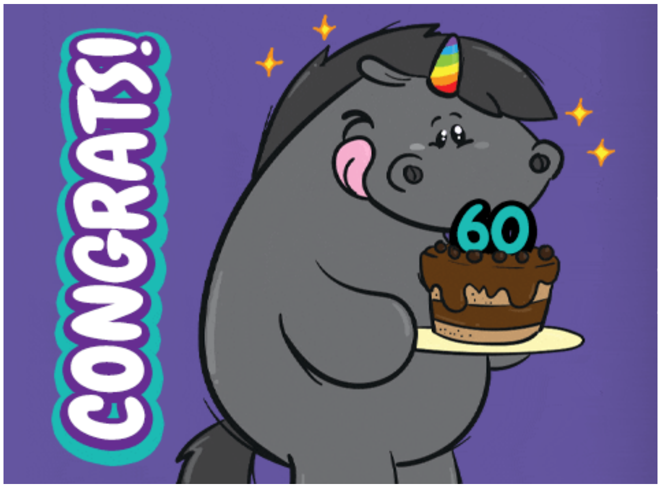 |

### Export

| | | |
|---|---|---|
| 1. | In the ‘ImageJ’ window:<br>a. Click ‘File’ → ‘Save As’ → ‘PNG…’<br>b. Include your name and ‘ImageJ’ in the file name<br>c. Click ‘Save’ | 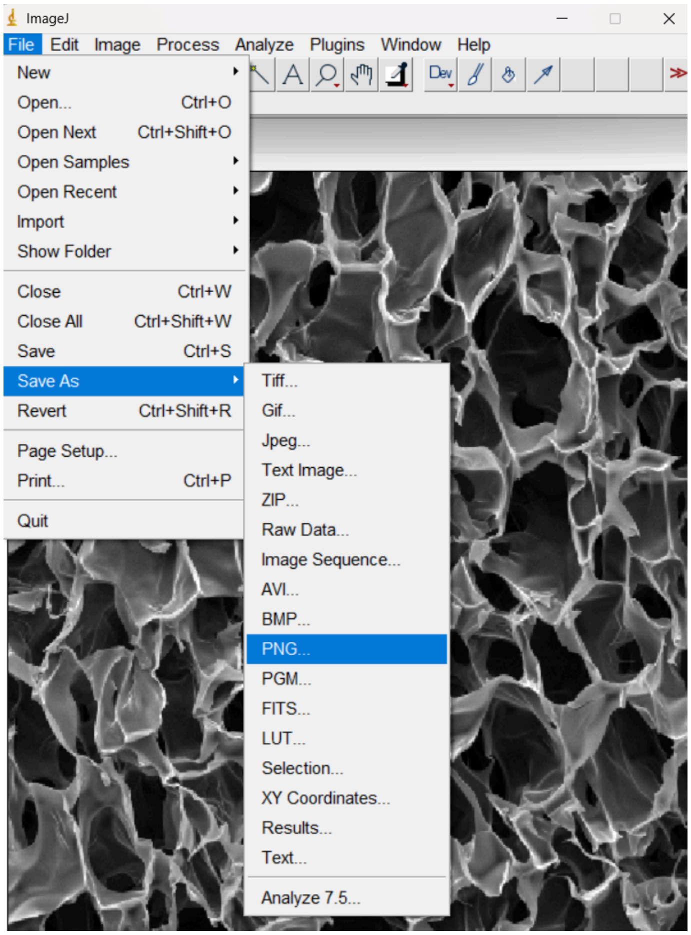 |

| | | |
|---|---|---|
| 2. | In the ‘Results’ window:<br>a. Click ‘File’ → ‘Save As…’<br>b. Include your name and ‘ImageJ’ in the file name<br>c. Click ‘Save’ | 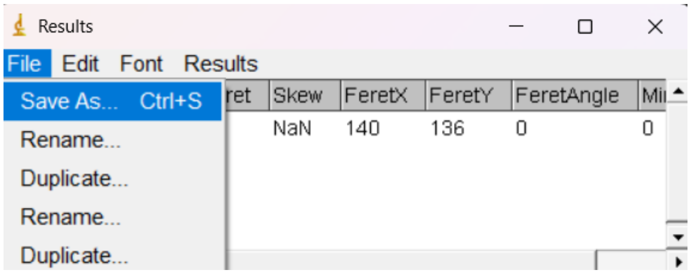 |
| 3. | Send Levi the two files by email or Slack | 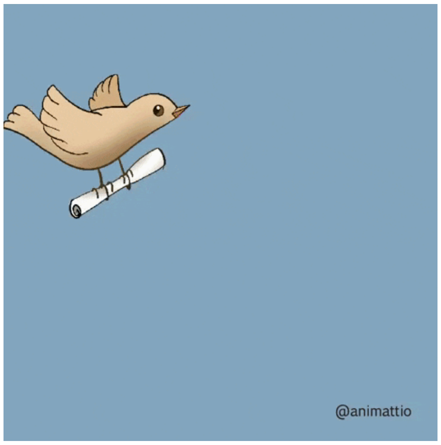 |
