## Supplementary material for "PoreVision: A Program for Enhancing Efficiency and Accuracy in SEM Pore Analyses": PoreVision Protocol.pdf

#### Tutorial

| | | |
|---|---|---|
| 1. | Open PoreVision | 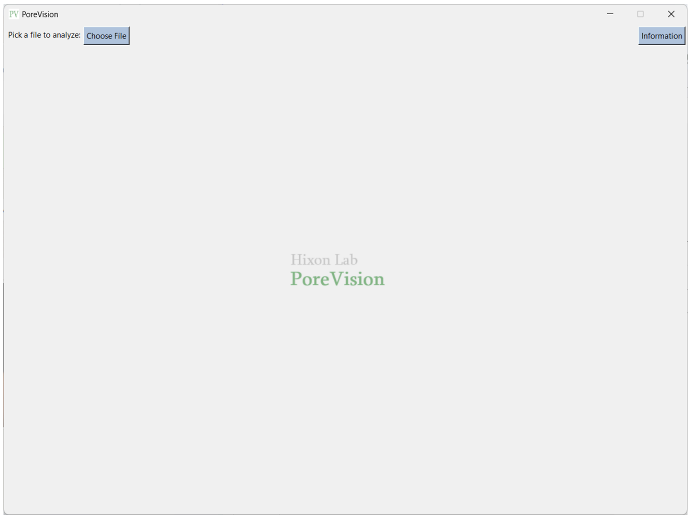 |
| 2. | Click ‘Choose File’ and select image | 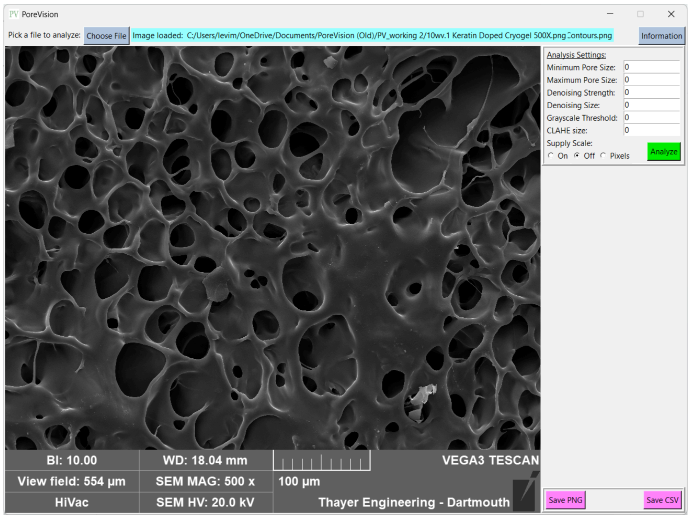 |

<table>
<tr>
<td>3.</td>
<td>

<u>Set the following values:</u><br>
Minimum Pore size: 10<br>
Maximum Pore size: 100000<br>
Denoising Strength: 0<br>
Denoising Size: 0<br>
Grayscale Threshold: 0<br>
CLAHE Size: 1<br>
<br>
Supply Scale: Off

</td>
<td>

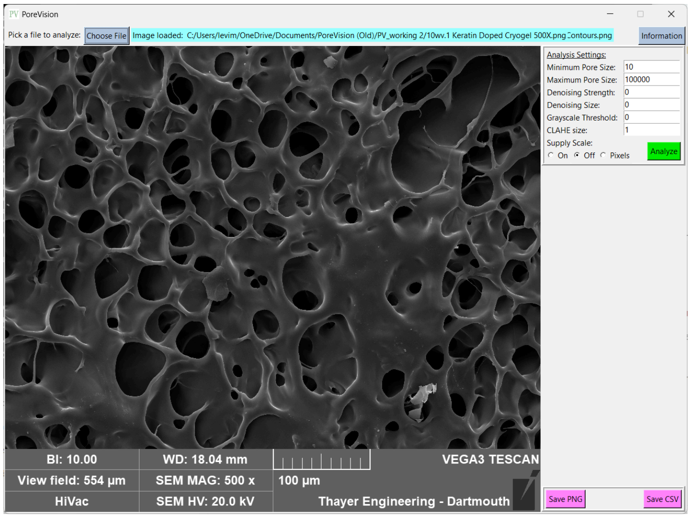


</td>
</tr>
<tr>
<td>4.</td>
<td>Click ‘Analyze’</td>
<td>

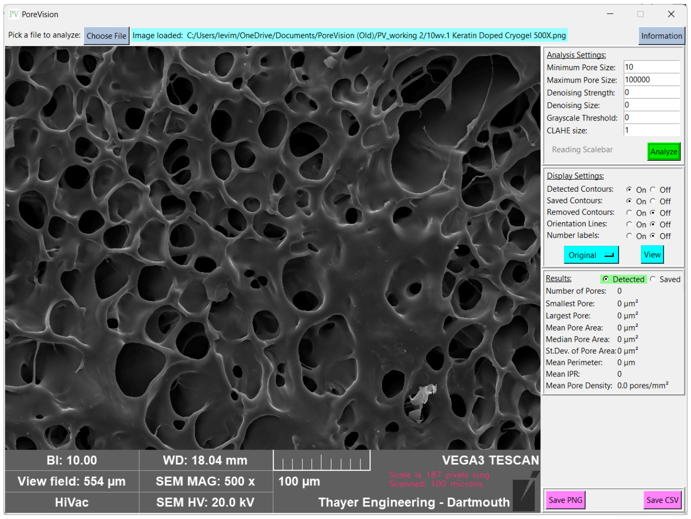


</td>
</tr>
<tr>
<td>5.</td>
<td>

Adjust each parameter under <u>Analysis Settings</u> to begin pore detection (I recommend to do this in the following order):<br>
<br>
Everytime you change a parameter, hit ‘Analyze’ to see how it affects the pore detection<br>
<br>
I. Adjust Grayscale Threshold<br>
A. Start in the middle of the recommended range (i.e. ~125)<br>
B. Click ‘Analyze’<br>
C. Are there too many outlines? Are there too few?<br>
D. Change the number and hit ‘Analyze’ again

</td>
<td>

<u>Recommended Parameter Ranges:</u><br>
<br>
Minimum Pore size: 3 to ∞<br>
Maximum Pore size: 3 to ∞<br>
Denoising Strength: 0 to 10000<br>
Denoising Size: 0 to 30<br>
Grayscale Threshold: 0 to 255<br>
CLAHE Size: 1 to 20

</td>
</tr>
</table>

E. Keep doing this until you are mostly satisfied with the pores detected
   a. You will be able to tune this and delete pores later so I recommend having more pores outlined than having fewer

II. Adjust **Minimum Pore Size** and **Maximum Pore Size**

A. If there are tiny outlines that seem like noise (e.g. dust or random particles)
   a. Increase **Minimum Pore Size** and hit 'Analyze'

B. If there are huge outlines that don't make sense
   a. Decrease **Maximum Pore Size** and hit 'Analyze'

III. Adjust **Denoising Strength**, **Denoising Size**, and **CLAHE Size**

A. These make small adjustments to the outlines (making them smoother and such)
   a. Change them in small increments to see how they affect the outlines

IV. You can click 'View' under Display Settings to view the image "under" the outlines

- This help see that the outlines are outlining the actual pores
- Click 'View' again to return the outlines

V. Repeat steps I-IV as needed until you are satisfied with the outlined pores (after this step you will be able to remove individual pores you don't like)

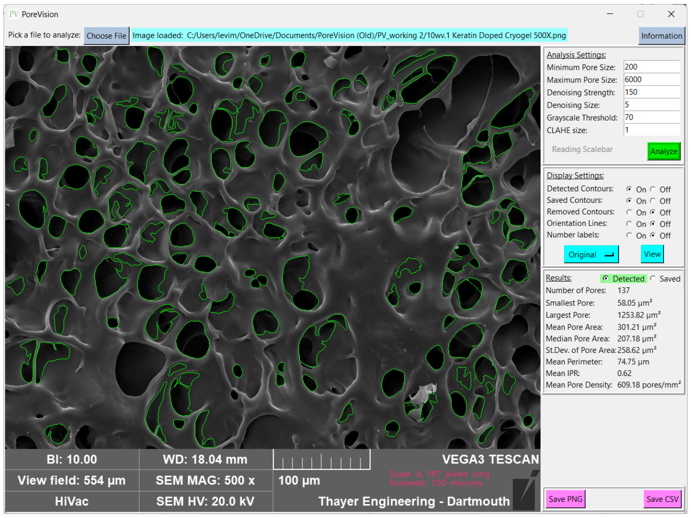

<table>
<tr>
<td>6.</td>
<td>Remove outlines you don’t like<br><br>Click on any outline and press ‘Remove’ in the bottom right to remove the outline<br><br>Tip: Click ‘View’ to confirm that the outline is erroneous; compare the pore underneath to the green outline</td>
<td>
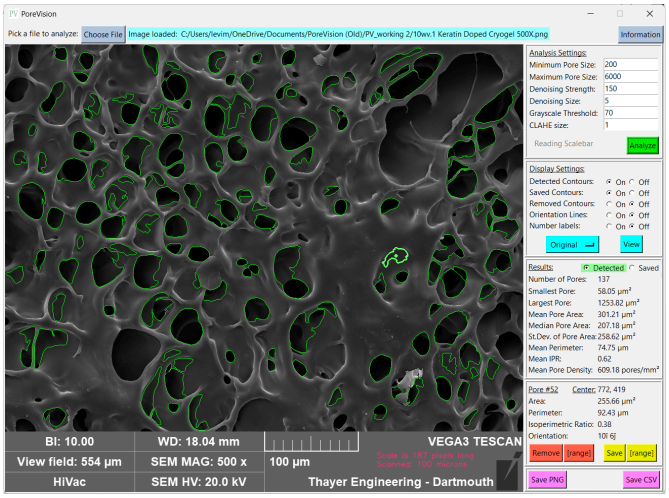

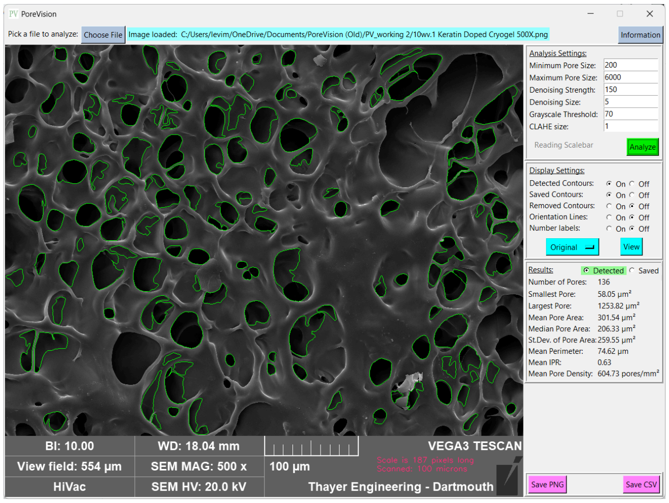

</td>
</tr>
<tr>
<td>7.</td>
<td>Save outlines you like<br><br>Click on any outline and press ‘Save’ in the bottom right to save the outline<br><br>Tip: Click ‘View’ to confirm that the outline is acceptable; compare the pore underneath to the green outline<br><br>Tip: Check the ‘On’ button next to ‘Saved Contours’ to hide the Saved pores; this can help reduce clutter on the screen</td>
<td>
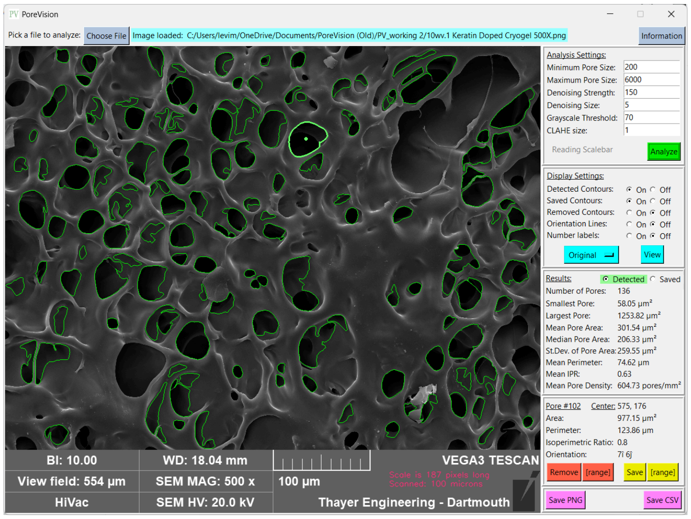

</td>
</tr>
</table>

<table>
<tr>
<td></td>
<td></td>
<td>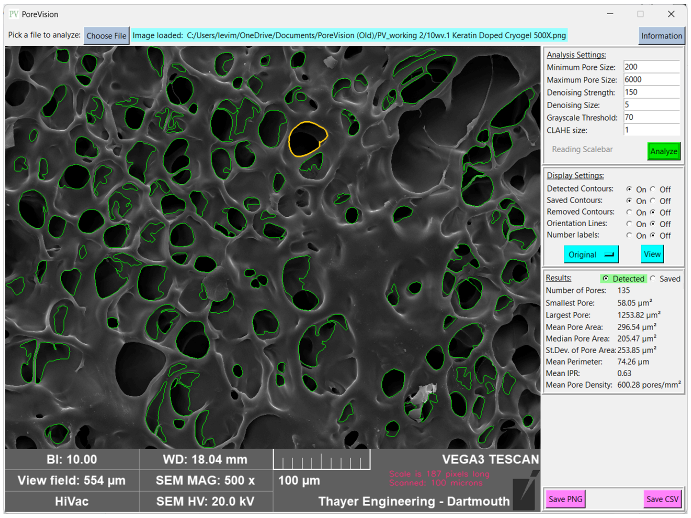
</td>
</tr>
<tr>
<td>8.</td>
<td>Save or remove multiple outlines<br><br>Check the ‘On’ button next to ‘Number labels’<br><br>Click on a pore and click [range] to remove a range of pores or [range] to save a range of pores</td>
<td>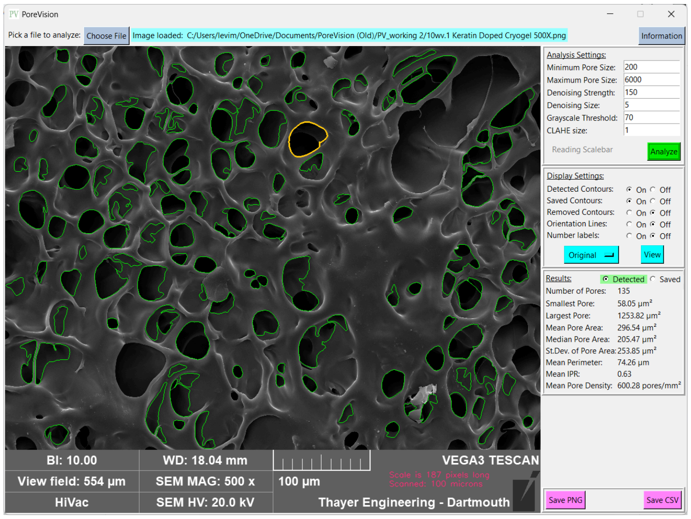

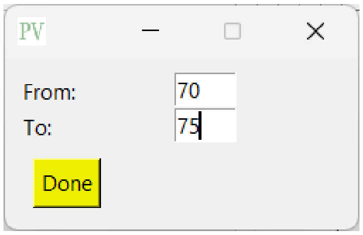
</td>
</tr>
</table>

| | | |
|---|---|---|
| | | 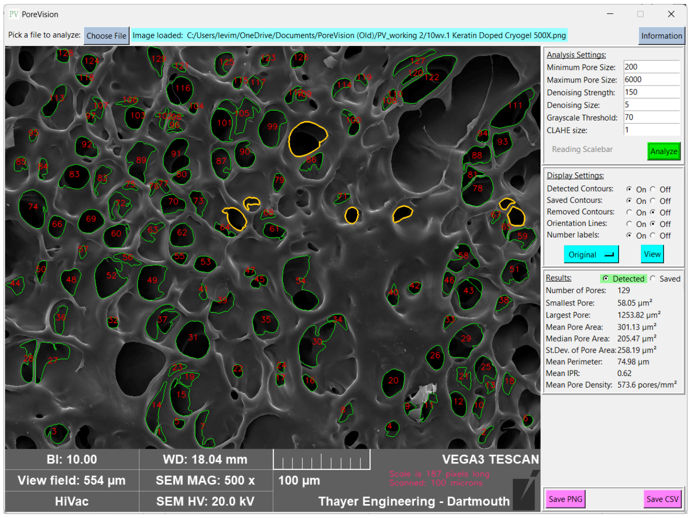 |
| 9. | Continue until all pores are either saved or removed<br><br>*Tip:*<br>*- Click 'Detected' next to 'Results,' there should be 0 for 'Number of Pores'*<br>*- Then click 'Saved' next to 'Results,' there should be a bunch of pores detected* | 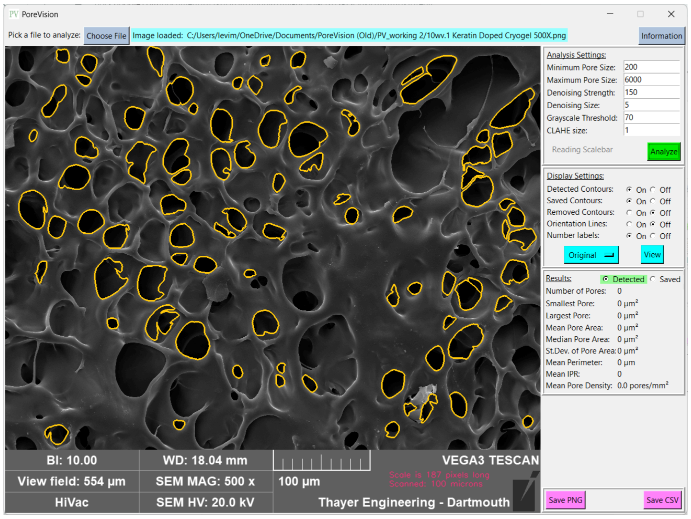<br>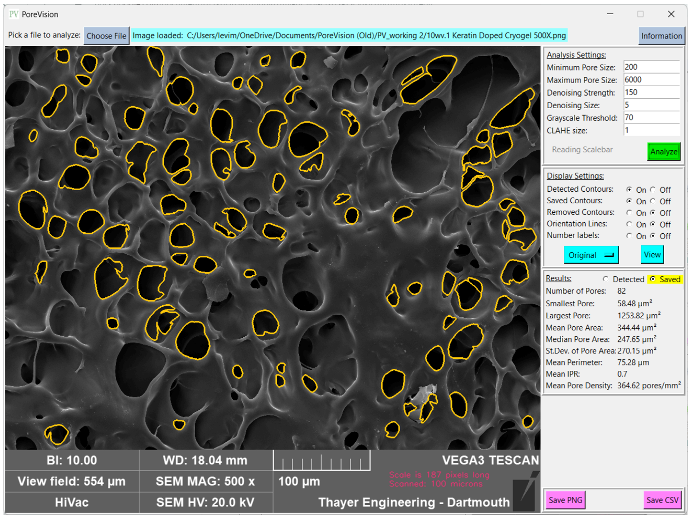 |

### Export

| | | |
|---|---|---|
| 1. | Under <u>Display Settings</u>, tick the following settings:<br>● **Detected Contours**: 'On'<br>● **Saved Contours**: 'On'<br>● **Removed Contours**: 'Off'<br>● **Orientation Lines**: 'Off'<br>● **Number labels**: 'On'<br><br>Next to <u>Results</u>:<br>● Tick the 'Saved' button | 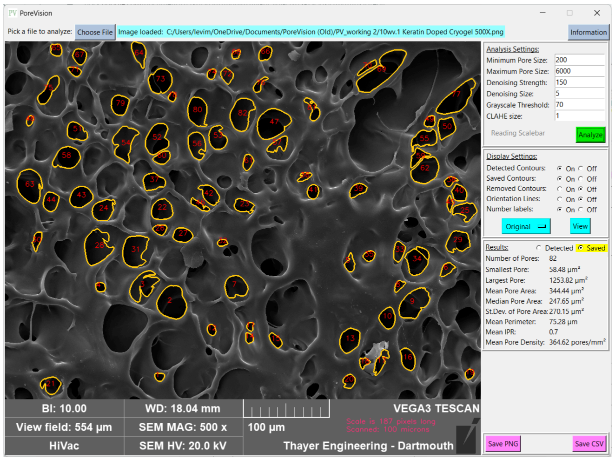 |
| 2. | Click 'Save PNG'<br><br>Include your name and 'PoreVision' in the file name | |
| 3. | Click 'Save CSV'<br><br>Include your name and 'PoreVision' in the file name | |
| 4. | Send Levi the two files by email or Slack | |
