## Supplementary figures and images for "PoreVision: A Program for Enhancing Efficiency and Accuracy in SEM Pore Analyses"

### Pore Measurement Test Image (Cropped).png

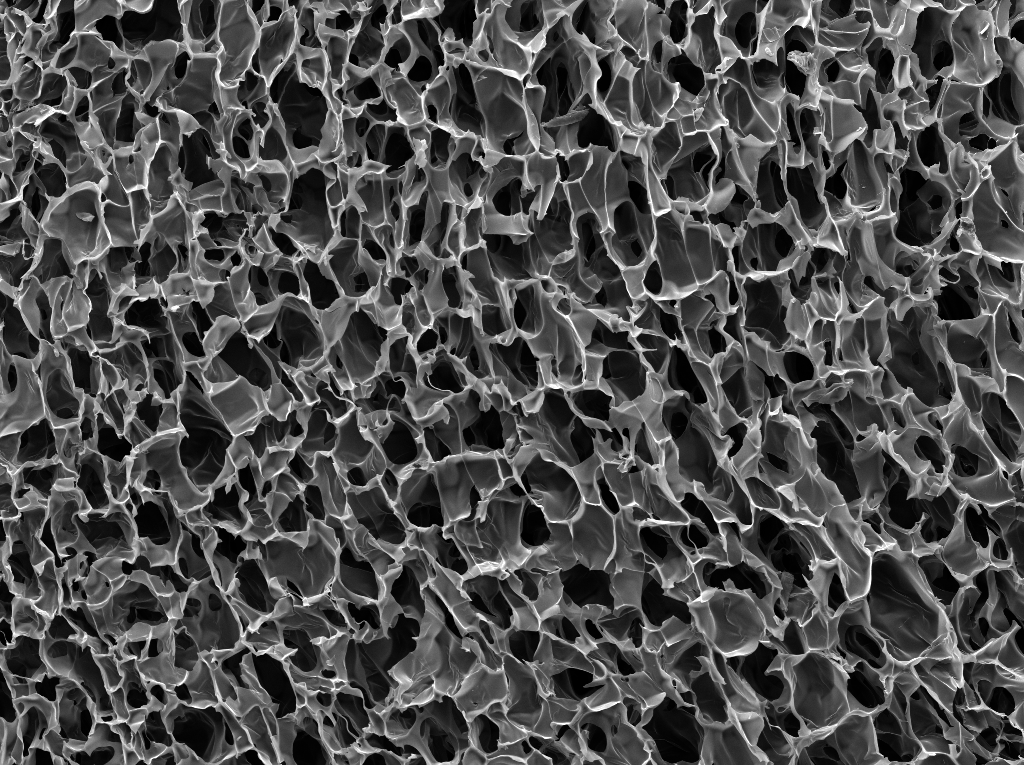

### Pore Measurement Test Image (Original).png

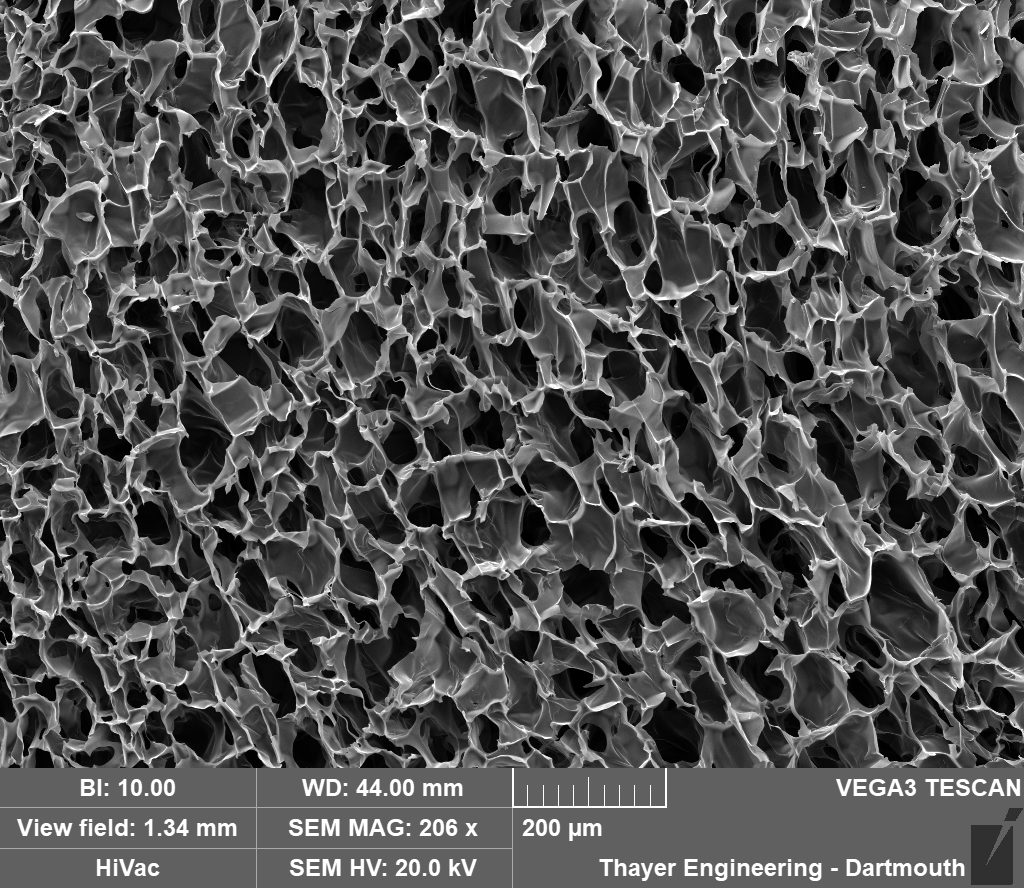

### Researcher 1 - ImageJ Image.png

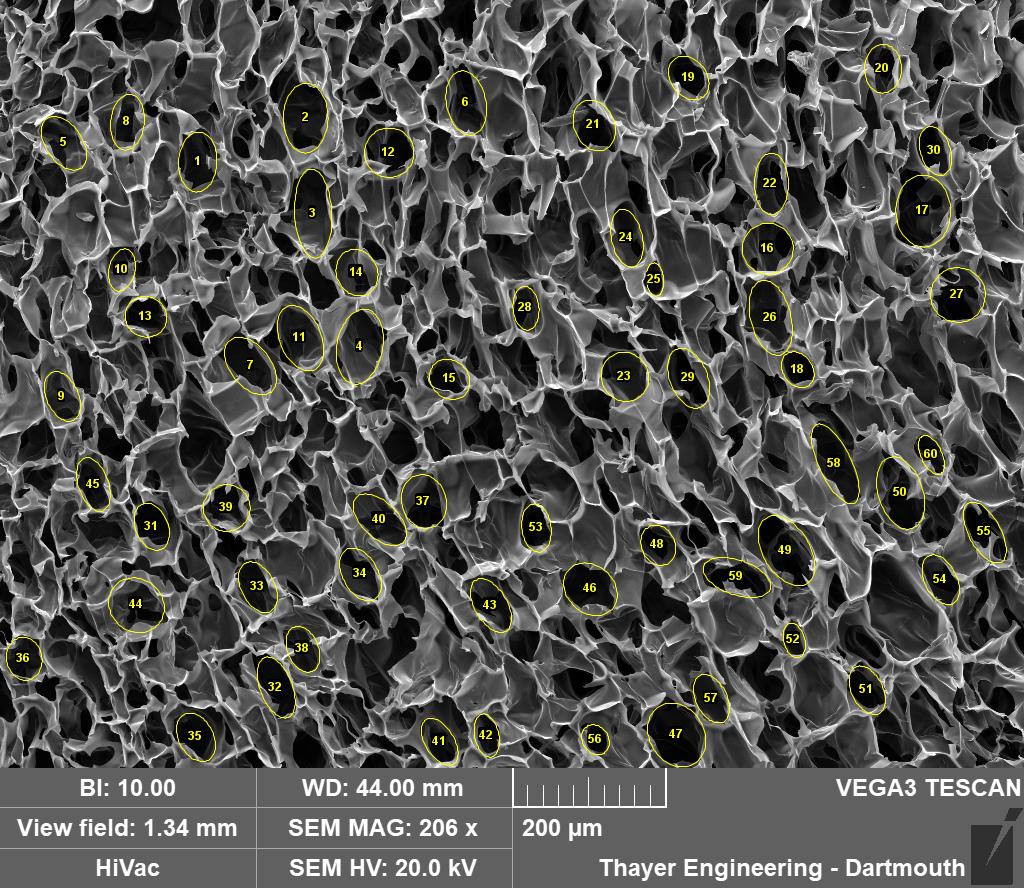

### Researcher 1 - PoreVision Image.png

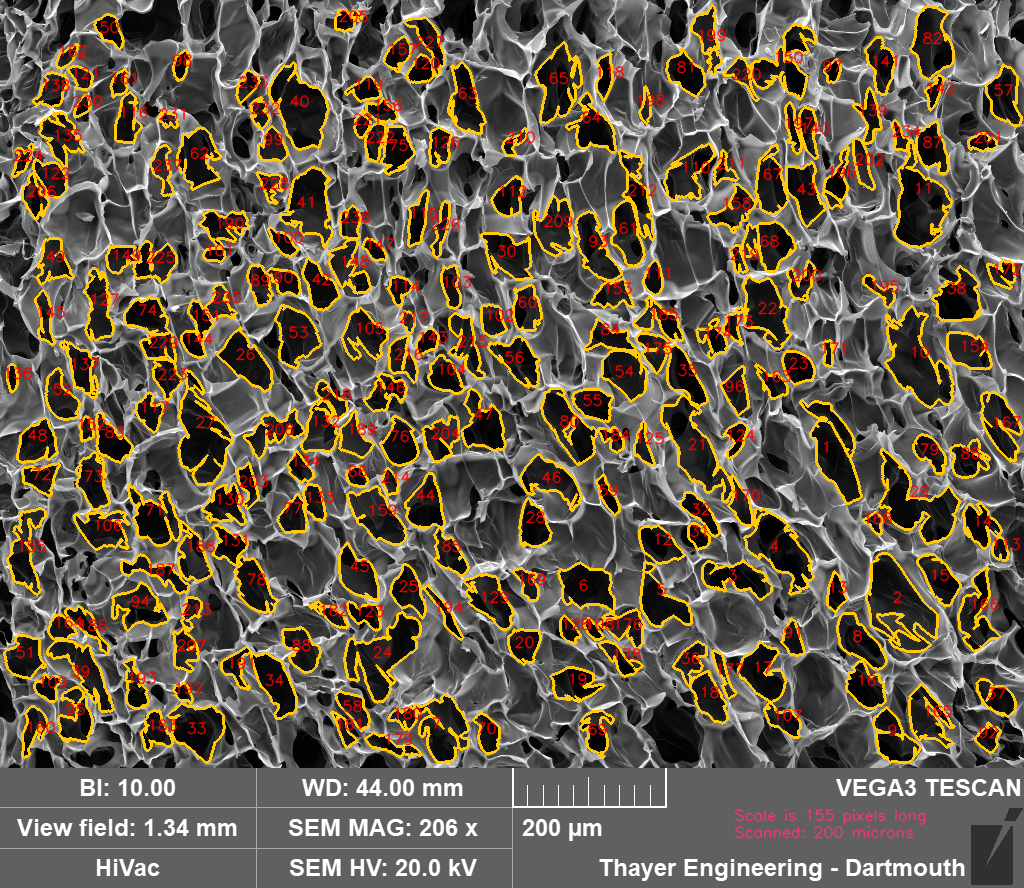

### Researcher 2 - ImageJ Image.png

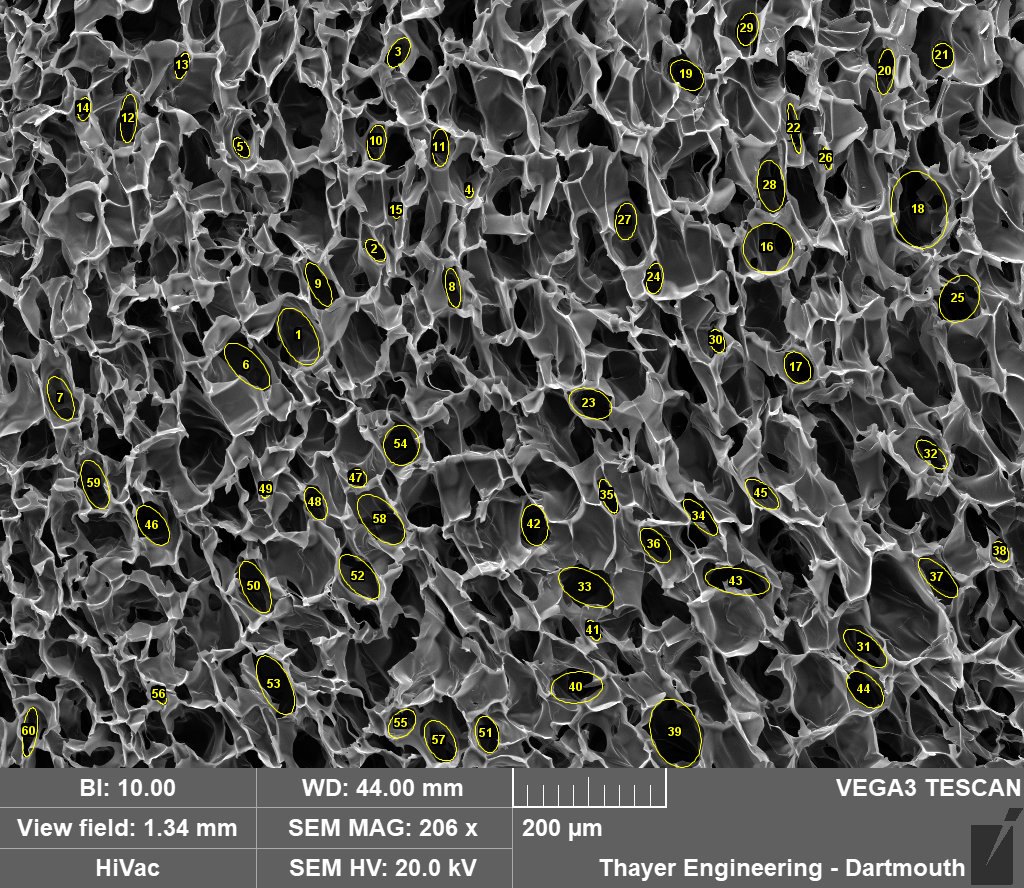

### Researcher 2 - PoreVision Image.png

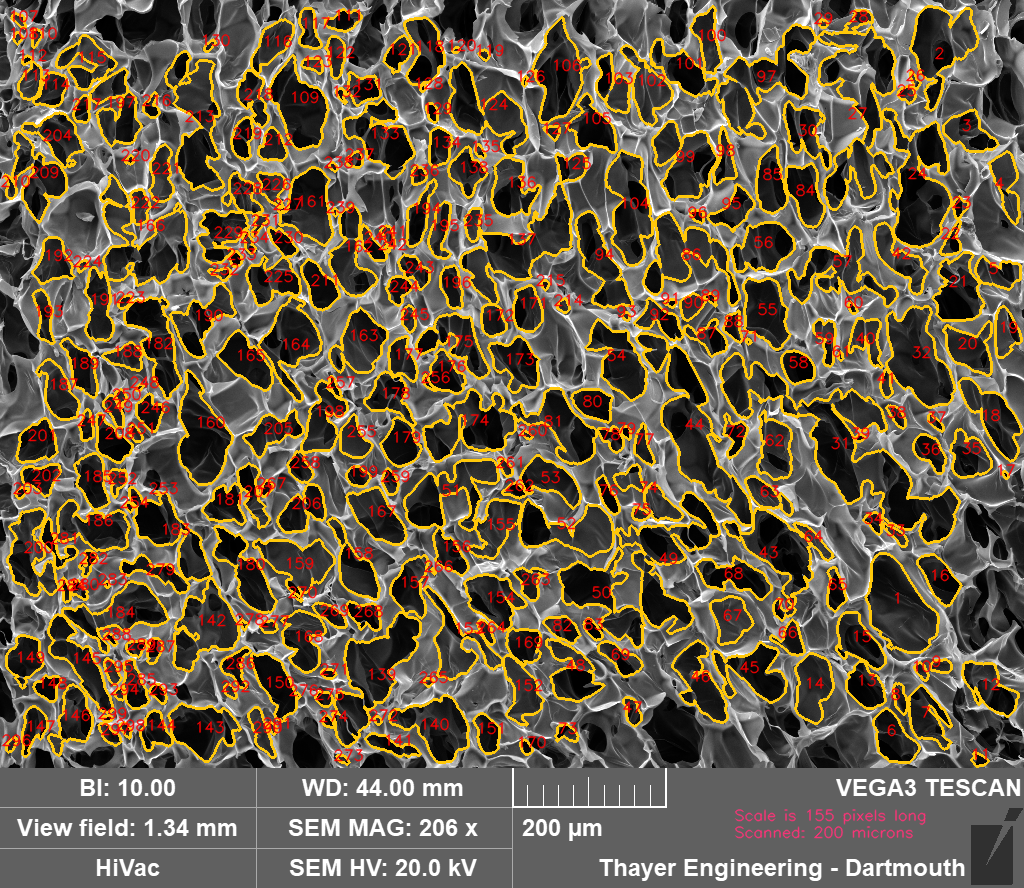

### Researcher 3 - ImageJ Image.png

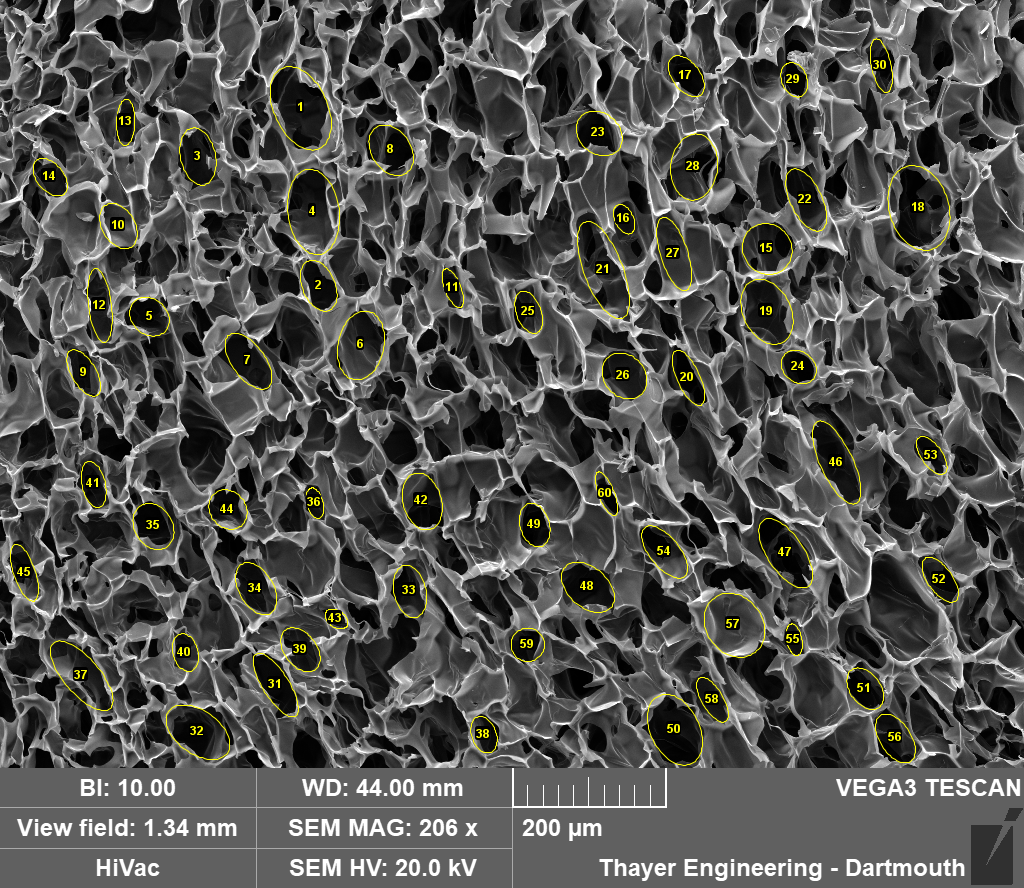

### Researcher 3 - PoreVision Image.png

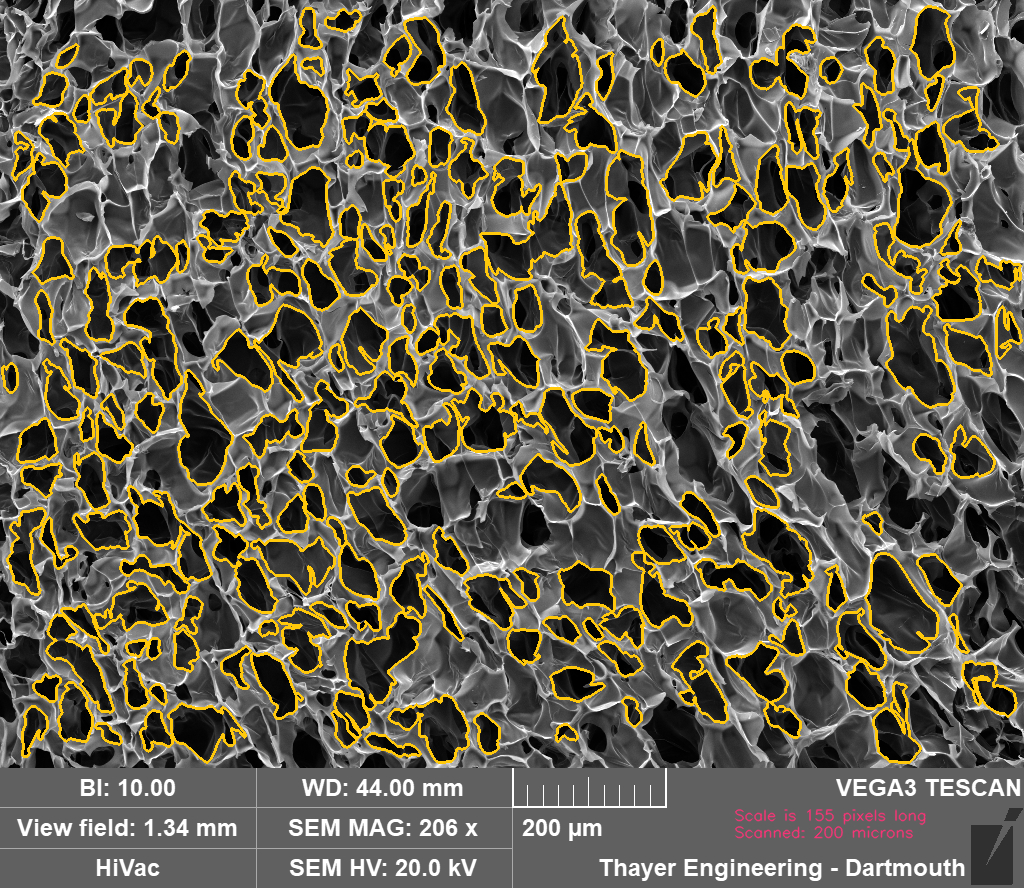

### Researcher 4 - ImageJ Image.jpg

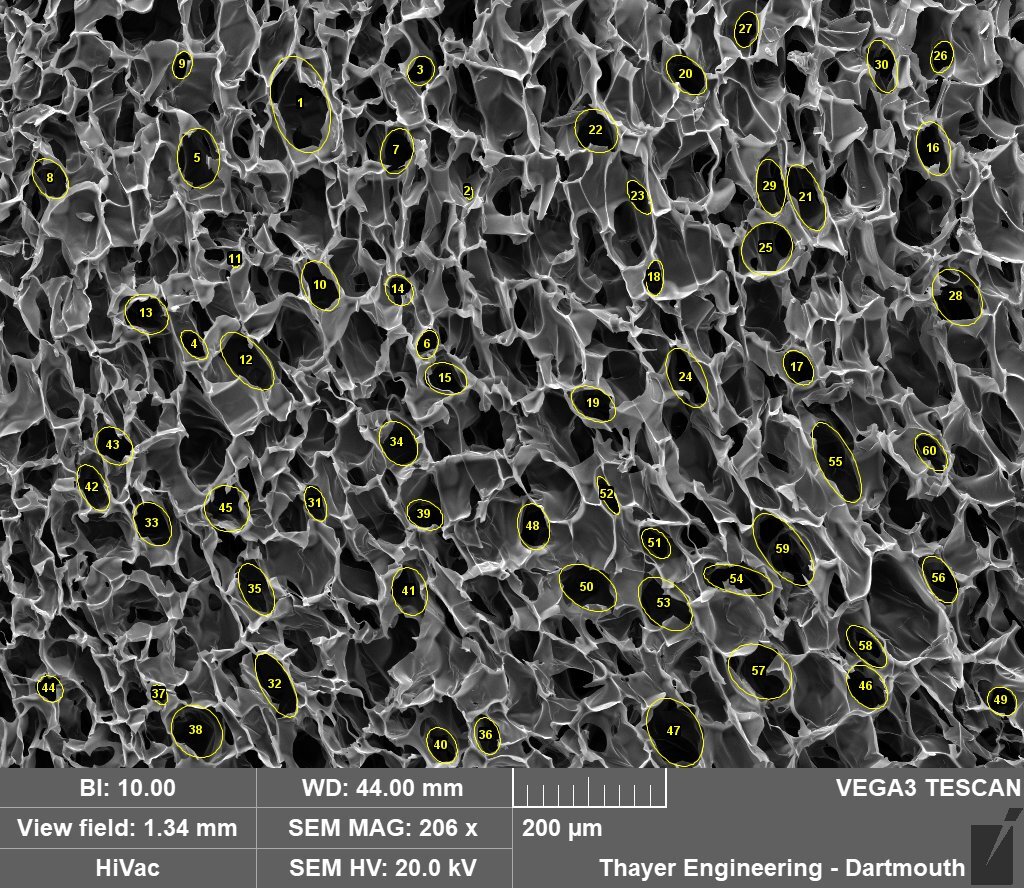

### Researcher 4 - PoreVision Image.png

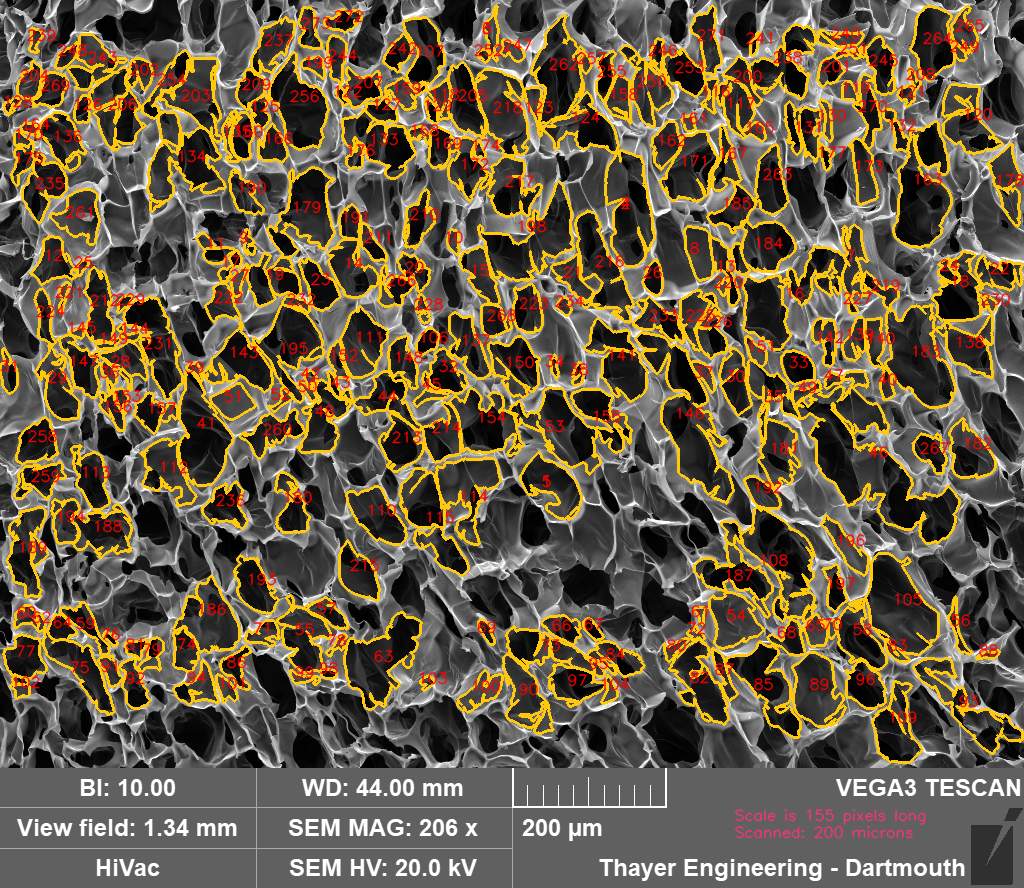

### Researcher 5 - ImageJ Image.png

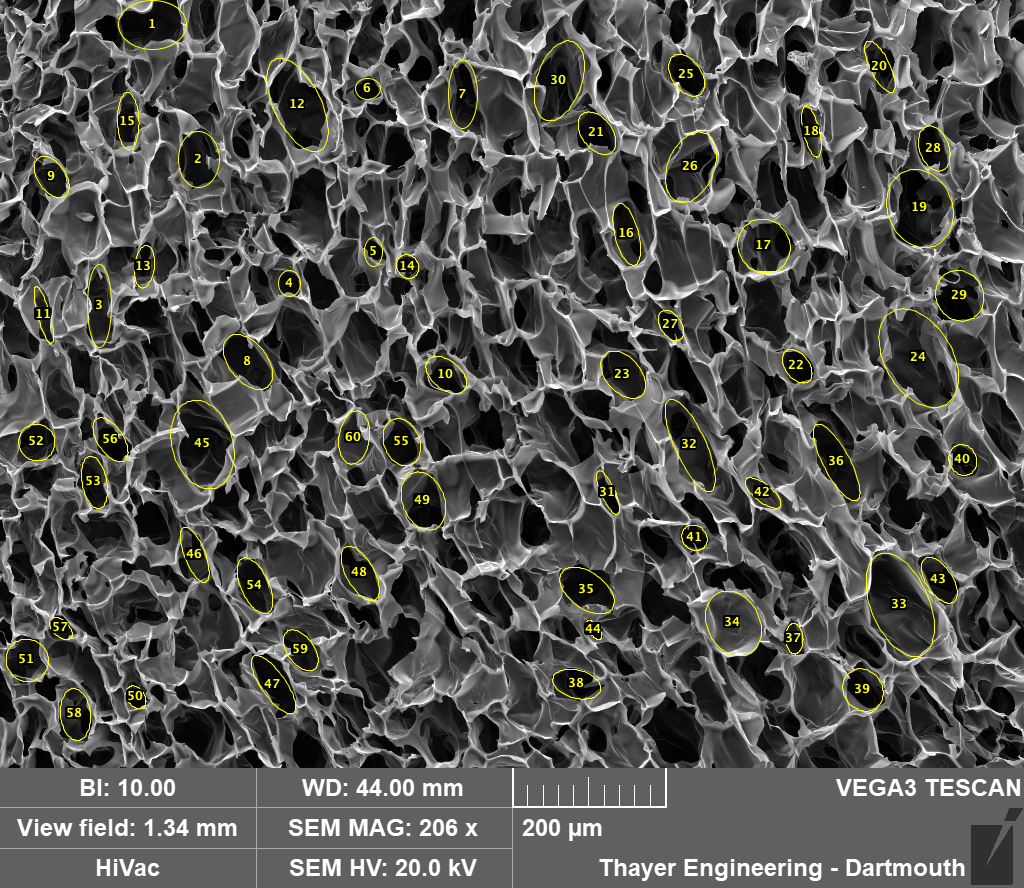

### Researcher 5 - PoreVision Image.png

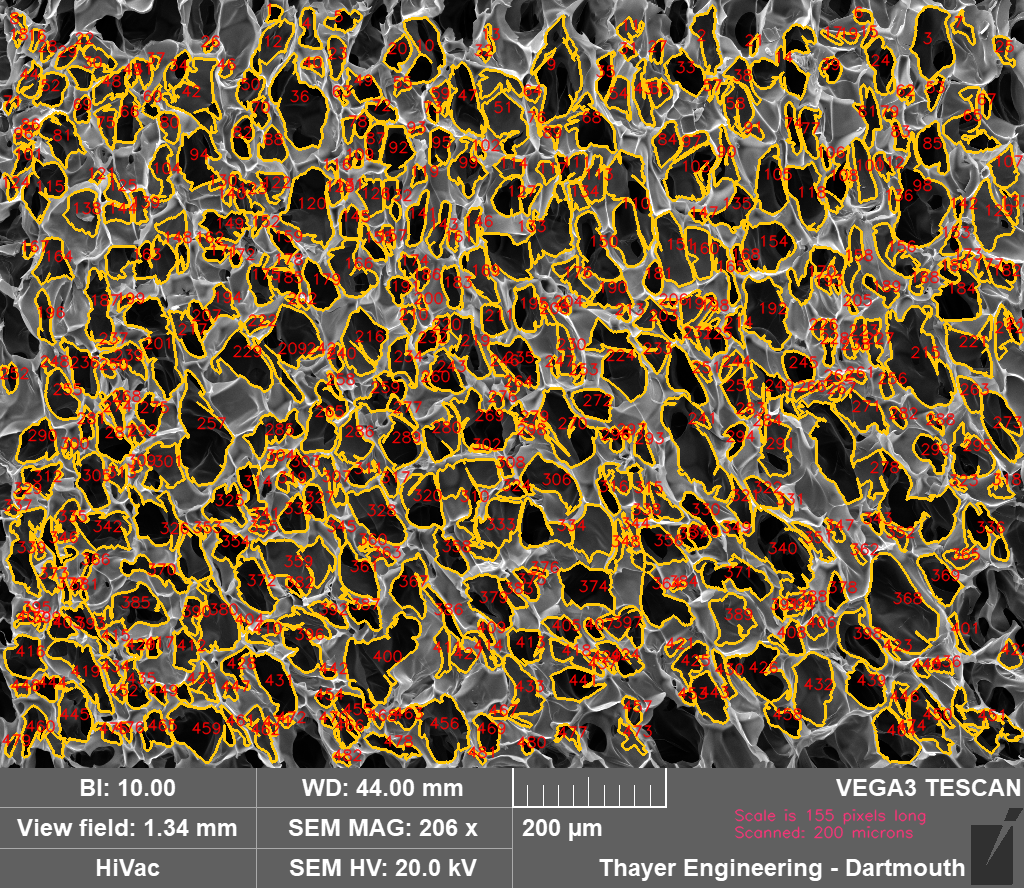
